# Conventional dose rate spatially-fractionated radiation therapy (SFRT) treatment response and its association with dosimetric parameters – A preclinical study in a Fisher 344 rat model

**DOI:** 10.1101/2020.01.30.926576

**Authors:** Judith N. Rivera, Thomas M. Kierski, Sandeep K. Kasoji, Anthony S. Abrantes, Paul A. Dayton, Sha X. Chang

## Abstract

**Purpose:** To identify key dosimetric parameters that have close associations with tumor treatment response and body weight change in SFRT treatments with a large range of spatial-fractionation scale at dose rates of several Gy/min.

**Methods:** Six study arms using uniform tumor radiation, half-tumor radiation, 2mm beam array radiation, 0.3mm minibeam radiation, and an untreated arm were used. All treatments were delivered on a 320kV x-ray irradiator. Forty-two female Fischer 344 rats with fibrosarcoma tumor allografts were used. Dosimetric parameters studied are peak dose and width, valley dose and width, peak-to-valley-dose-ratio, volumetric average dose, percentage volume directly irradiated, and tumor- and normal-tissue EUD. Animal survival, tumor volume change, and body weight change (indicative of treatment toxicity) are tested for association with the dosimetric parameters using linear regression and Cox Proportional Hazards models.

**Results:** The dosimetric parameters most closely associated with tumor response are tumor EUD (R^2^=0.7923, F-stat=15.26*; z-test=−4.07***), valley/minimum dose (R^2^=0.7636, F-stat=12.92*; z-test=−4.338***), and percentage tumor directly irradiated (R^2^=0.7153, F-stat=10.05*; z-test=−3.837***) per the linear regression and Cox Proportional Hazards models, respectively. Tumor response is linearly proportional to valley/minimum doses and tumor EUD. Average dose (R^2^=0.2745, F-stat=1.514 (no sig.); z-test=−2.811**) and peak dose (R^2^=0.04472, F-stat=0.6874 (not sig.); z-test=−0.786 (not sig.)) show the weakest associations to tumor response. Only the uniform radiation arm did not gain body weight post-radiation, indicative of treatment toxicity; however, body weight change in general shows weak association with all dosimetric parameters except for valley/min dose (R^2^=0.3814, F-stat=13.56**), valley width (R^2^=0.2853, F-stat=8.783**), and peak width (R^2^=0.2759, F-stat=8.382**).

**Conclusions:** For a single-fraction SFRT at conventional dose rates, valley, not peak, dose is closely associated with tumor treatment response and thus should be used for treatment prescription. Tumor EUD, valley/min dose, and percentage tumor directly irradiated are the top three dosimetric parameters that exhibited close associations with tumor response.

## Introduction

Spatially-fractionated radiation therapy (SFRT) is a nonconventional radiation therapy that is characterized by intentionally-created high dose inhomogeneities, ultra-high maximum doses, and single fraction treatments (*1, 2*). The dose inhomogeneity consists of many small sub-regions with alternating high and low doses throughout the treatment volume. SFRT includes clinical GRID therapy(*1, 3*) and preclinical microbeam radiation therapy (MRT) (*4*), each of which of has a decades-long history demonstrating its superior therapeutic ratio compared to conventional radiation therapy, especially in terms of normal organ sparing. Detailed summaries can be found in two recent reviews by Billena and Khan (*5*) for GRID therapy and by Eling et al. (6, 7) for MRT. Today, there are a number of modern treatment delivery technologies available for clinical SFRT including multi-leaf collimator generated GRID (*8*), LATTICE (*9–11*), Tomotherapy (*5*), and particle GRID therapy (*12, 13*). For preclinical SFRT, newer technologies include “minibeams” with larger spatial fractionation scales (on the order of millimeter instead of the tens of microns used in classical MRT) (*14, 15*) and with conventional dose-rates (*16, 17*). Most published MRT research utilized brilliant x-rays generated from synchrotron accelerator facilities with ultrahigh dose rates (*4*). The conventional dose rate SFRT radiations, such as the ones used in this study, are highly relevant to translational research for LINAC-based SFRT clinical applications, where conventional dose rates are also used.

Despite the long history and well demonstrated therapeutic ratio advantage over conventional uniform dose radiation therapy, SFRT remains an experimental therapy. There are several reasons attributed to the sluggish clinical translation progress including a lack of understanding of SFRT working mechanisms and of the association between SFRT treatment response and dosimetry. While we have verified treatment dosimetry and tumor control outcome correlations for conventional radiation therapy (i.e., tumor minimum dose and EUD are closely correlated with tumor control) (*18*) we do not yet have such understanding for SFRT, which has significantly more complex dosimetry than that of conventional radiation therapy. Unique SFRT dosimetric parameters that describe the dosimetry include peak dose, valley dose, peak-to-valley-dose-ratio, peak width, valley width, and percentage tumor volume directly irradiated. It is reasonable to assume that not all these dosimetric parameters have the same clinical significance. To effectively advance SFRT clinical translation it is critically important to identify which parameters have strong/weak associations with a given treatment response.

The goal of this study is to identify key dosimetric parameters that are most closely associated with treatment response using a preclinical animal model. We hypothesize that while peak dose has always been used to prescribe SFRT treatment for both clinical and preclinical applications, peak dose may not be the dosimetric parameter most closely associated with SFRT tumor control or treatment toxicity. If it is not, which SFRT dosimetric parameters are? Further, we ask that, for a given pattern of SFRT treatment, what is its conventional radiation therapy equivalence for a given treatment response? The answers to these questions are crucial to advance clinical translation of SFRT. Unfortunately, decades of synchrotron-based MRT studies may not be able to answer these questions due to the use of ultrahigh dose rates (1000sGy/sec) (*19*). Recent research on FLASH radiation has shown that radiation with dose-rates of 100Gy/s or higher selectively spares normal tissue not tumor (*4, 20, 21*). This new finding revealed that the ultrahigh dose-rate alone is partially responsible for the observed high therapeutic-ratio demonstrated in the majority of SFRT research published so far (6). This study will help discern the impact of radiation spatial fractionation at dose rates relevant to clinical SFRT treatments.

Today, SFRT is receiving much deserved renewed attention and enthusiasm in the field of radiation oncology. In 2018 National Cancer Institute and Radiosurgery Society jointly held the first workshop on Understanding High-Dose, Ultra-Dose-Rate and Spatially Fractionated Radiotherapy and created three standing working groups (clinical, biology, and physics) aiming to provide guidelines on SFRT research and clinical application (*22*). We hope this work will assist in this endeavor by shedding light on the clinical impact of SFRT dosimetry parameters.

## Materials and methods

### Study design

The secret of SFRT lays in its radiation dose spatial fractionation. Although this work does not address the very much needed understanding of working mechanism it addresses another important matter for SFRT application - the association of SFRT dosimetric parameters with treatment response at conventional dose rates (dose rate ranges from 4.27 to 5.25Gy/min was used). Fig 1 shows a six-arm study design using a very large span of radiation spatial fractionation, constructed to explore the impact of radiation spatial fractionation. Table 1 summarizes the dosimetric parameters of each of the six arms. To study the effect of radiation spatial fractionation under the condition of equal volume-averaged dose we used the following four study arms: 20GyUniformRT (entire tumor directly irradiated), 20GyHalfRT (only one-half of tumor directly irradiated), 20Gy2mmSFRT (50% of tumor directly irradiated by 2mm-wide planar beam array), and 20GySFRT (20% of tumor directly irradiated with 0.3mm-wide planar beam array). Note that the doses are volume-averaged doses computed for the entire tumor volume. A 50GySFRT arm (50Gy volume-averaged dose, beam width 0.31mm) is added as it has a peak dose of 225Gy, which is within the known minibeam peak dose range showing tumor control. To account for unavoidable variations in tumor position under the 20Gy2mmSFRT treatment beams during animal irradiations, we computed the maximum and minimum beam coverage positions and calculated their corresponding dosimetric specifications. The 20Gy2mmSFRT treatment arm dosimetric values reported in Table 1 correspond to the average at these positions for a 10mm diameter tumor. For example, a 10mm sized tumor is irradiated by at most three 2mm-peaks and at minimum two 2mm-peaks and the average dosimetric parameter at these two positions was calculated.

**Fig 1.**
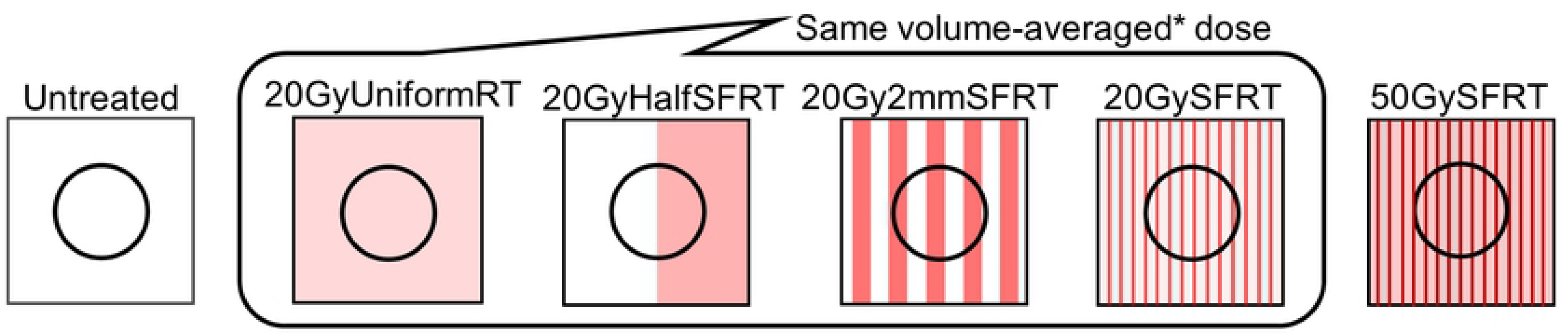
Illustration of the SFRT spatial fractionation study design. A very large range of radiation spatial fractionation scale was used to derive the impact of radiation spatial fractionation. Four arms share the same 20Gy volume-average dose. The high dose 50GySFRT arm is added because 20GySFRT is not known to have tumor control. The dosimetric parameters studied and number of animals per study arm are listed in Table 1.

**Table 1.**
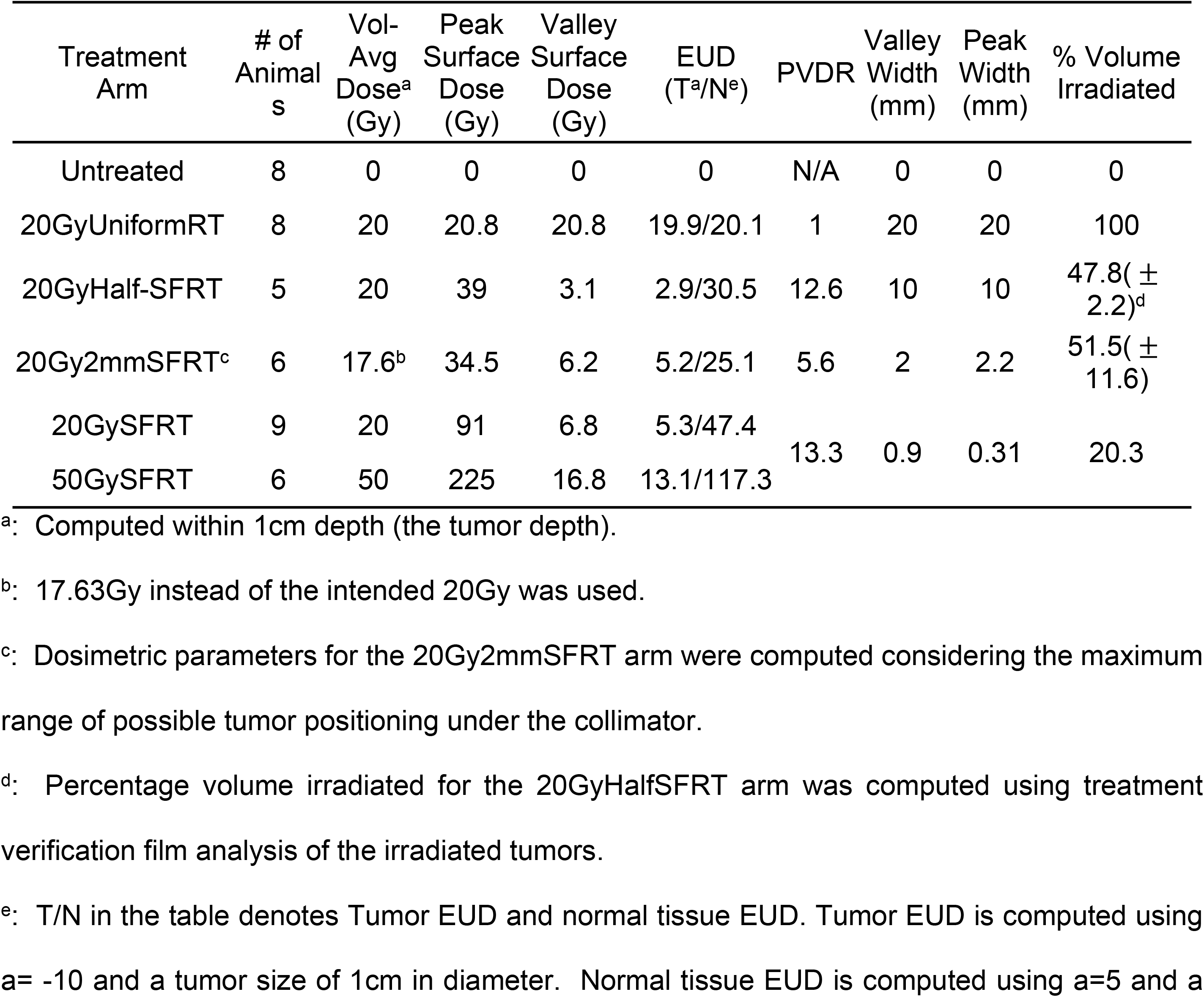
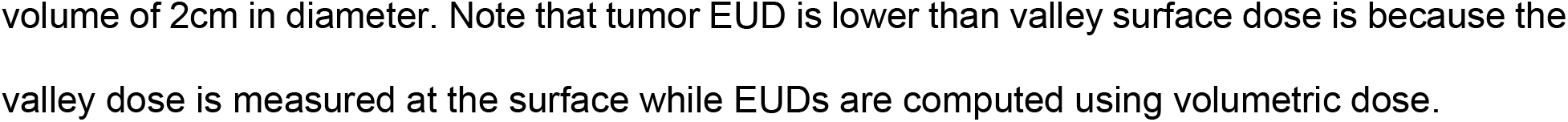
Summary of nine SFRT dosimetric parameter specifications in the six-arm study.

| Treatment Arm | # of Animals | Vol-Avg Dose<sup>a</sup> (Gy) | Peak Surface Dose (Gy) | Valley Surface Dose (Gy) | EUD (T<sup>a</sup>/N<sup>e</sup>) | PVDR | Valley Width (mm) | Peak Width (mm) | % Volume Irradiated |
| --- | --- | --- | --- | --- | --- | --- | --- | --- | --- |
| Untreated | 8 | 0 | 0 | 0 | 0 | N/A | 0 | 0 | 0 |
| 20GyUniformRT | 8 | 20 | 20.8 | 20.8 | 19.9/20.1 | 1 | 20 | 20 | 100 |
| 20GyHalf-SFRT | 5 | 20 | 39 | 3.1 | 2.9/30.5 | 12.6 | 10 | 10 | 47.8( ± 2.2)<sup>d</sup> |
| 20Gy2mmSFRT<sup>c</sup> | 6 | 17.6<sup>b</sup> | 34.5 | 6.2 | 5.2/25.1 | 5.6 | 2 | 2.2 | 51.5( ± 11.6) |
| 20GySFRT | 9 | 20 | 91 | 6.8 | 5.3/47.4 | 13.3 | 0.9 | 0.31 | 20.3 |
| 50GySFRT | 6 | 50 | 225 | 16.8 | 13.1/117.3 |  |  |  |  |

<sup>a</sup>: Computed within 1cm depth (the tumor depth).
<sup>b</sup>: 17.63Gy instead of the intended 20Gy was used.
<sup>c</sup>: Dosimetric parameters for the 20Gy2mmSFRT arm were computed considering the maximum range of possible tumor positioning under the collimator.
<sup>d</sup>: Percentage volume irradiated for the 20GyHalfSFRT arm was computed using treatment verification film analysis of the irradiated tumors.
<sup>e</sup>: T/N in the table denotes Tumor EUD and normal tissue EUD. Tumor EUD is computed using a= -10 and a tumor size of 1cm in diameter. Normal tissue EUD is computed using a=5 and a
volume of 2cm in diameter. Note that tumor EUD is lower than valley surface dose is because the valley dose is measured at the surface while EUDs are computed using volumetric dose.

Custom-made radiation blocks and collimators made of Cerrobend or tungsten were used to define the 2cm×2cm field for 20GyUniformRT arm treatment, the 2cm×1cm for 20GyHalfSFRT treatment, and the beamlet array 2cm×2cm fields for both the 20Gy2mmSFRT and 20Gy/50GySFRT treatments. The 2cm field size in the direction of the uniform dose within each of SFRT planar beams is made possible by the very large focal spot size (8mm^2^) of the XRad irradiator (Precision X-ray Inc., North Branford, CT USA). All irradiations in this study used the same irradiator.

### Animal tumor model

This study was carried out in strict accordance with the recommendations in the Guide for the Care and Use of Laboratory Animals of the National Institutes of Health (NIH). The University of North Carolina- Chapel Hill Institutional Animal Care and Use Committee (IACUC) reviewed and approved the animal protocol (IACUC ID: 15-366.0) in accordance with NIH standards. All animal surgical, radiation, and imaging procedures were performed under general anesthesia and all efforts were made to minimize suffering.

Forty-two eight-week-old female Fischer 344 rats from Charles River Labs and rat fibrosarcoma tumor allografts were used (*23*). The rat fibrosarcoma (FSA) allograft model has been well characterized in several radiotherapy response studies by our and collaborator labs (*23–25*). Rat FSA is characterized as a local, non-metastasizing tumor that is highly vascular and oxygen dependent (*34,35*). It is an appropriate tumor model for our long-term study goal that investigates the association of SFRT dosimetric parameters with treatment responses, which is reported here, and the association between SFRT treatment response and tumor vascular change post radiation using 3D acoustic angiography. The latter is ongoing research for future publication.

All surgical, radiation, and imaging procedures were performed under general anesthesia, induced in the animals initially using 5% vaporized isoflurane mixed with pure oxygen as the carrier gas and then maintained at 2.5% isoflurane mixed with pure oxygen throughout each procedure. Depth of anesthesia was monitored by toe pinch reflex and breathing rate. Opthalmic ointment was placed on the animal’s eyes during anesthesia to provide lubrication and body temperature under anesthesia was maintained via electronically controlled heating pad. Tumors were grown in each rat by implanting freshly resected tumor tissue (1mm^3^) that was harvested from tumor-bearing donor rats into the subcutaneous space of the rodent flank using blunt dissection. Postoperative care included daily incision surveillance, body temperature monitoring, and a water bottle containing 6mg/mL cherry-flavored, dye-free children’s Tylenol diluted in water for a minimum of 24-hrs post-surgery to alleviate any associated pain from the implantation procedure. Animals were used for experiments 2-3 weeks post-implantation, when the tumors reached the target RT treatment size of approximately 5-10mm.

In preclinical studies the pre-treatment tumor volume is known to be strongly correlated with treatment tumor control (*23*). We minimize this unwanted effect by controlling the pre-treatment tumor volume in a randomized, matched group study design. We binned animals according to their pre-treatment tumor volume and then randomly assigned these matched bins of animals such that at least one animal from each bin is assigned to each treatment group. This technique resulted in an average initial tumor volume across groups of 566 +/− 47 mm^3^ on RT treatment day. Biological variability was minimized by ordering animals from the same vendor and of the same age (6 weeks old), implanting tumor on the same day and from the same donor animal, treating with radiation on the same day, and housing animals in the same Vivarium location with identical husbandry conditions. All animals (mixed caged) were provided identical standard laboratory rodent diets of (23%> crude protein) and water ad libitum throughout the study. In addition, all animal diets were supplemented with high-calorie, nutritionally fortified water-based gel cups to help mitigate any potential significant weight loss and dehydration post-radiation.

The animals body weight and tumor volumes are monitored prior to radiation and every third day thereafter for up to 30 days. Study endpoints are maximum tumor burden (2.5cm or larger in any dimension), weight loss in excess of 15%, body condition scores (26) less than or equal to 2, or other signs of pain, discomfort, or moribundity as recommended by University of North Carolina- Chapel Hill Division of Comparative Medicine veterinary staff. Animals that met study end-point criteria will be ethically euthanized primarily via compressed carbon dioxide gas or vaporized isoflurane overdose followed by thoracotomy as a secondary means of physical euthanasia per the approved animal study protocol.

### Animal radiation dosimetry

XRad Irradiator and 320kV x-rays were used in this study. Surface dose rates ranging from 4.27 to 5.25Gy/min were used for all study arms. Fig 2 shows the treatment setup, the radiation light field on animal seen by the camera, and treatment verification films. Dosimetry was measured via EBT-3 film calibrated by an ADCL-calibrated ion chamber under large field conditions. Acrylic phantom measurement setup and beam profile and percentage depth dose (PDD) dosimetry are shown in Figs 3 and 4.The volume-averaged tumor dose was approximated by computing the film average dose within an area of 1cm by 1cm (depth) of the PDD film. The differential dose volume histograms of the PDD films were used for tumor and normal tissue EUD calculations as described by Niemierko (27) using values of a=−10 for tumor and a=5 for normal tissue.

**Fig 2.**
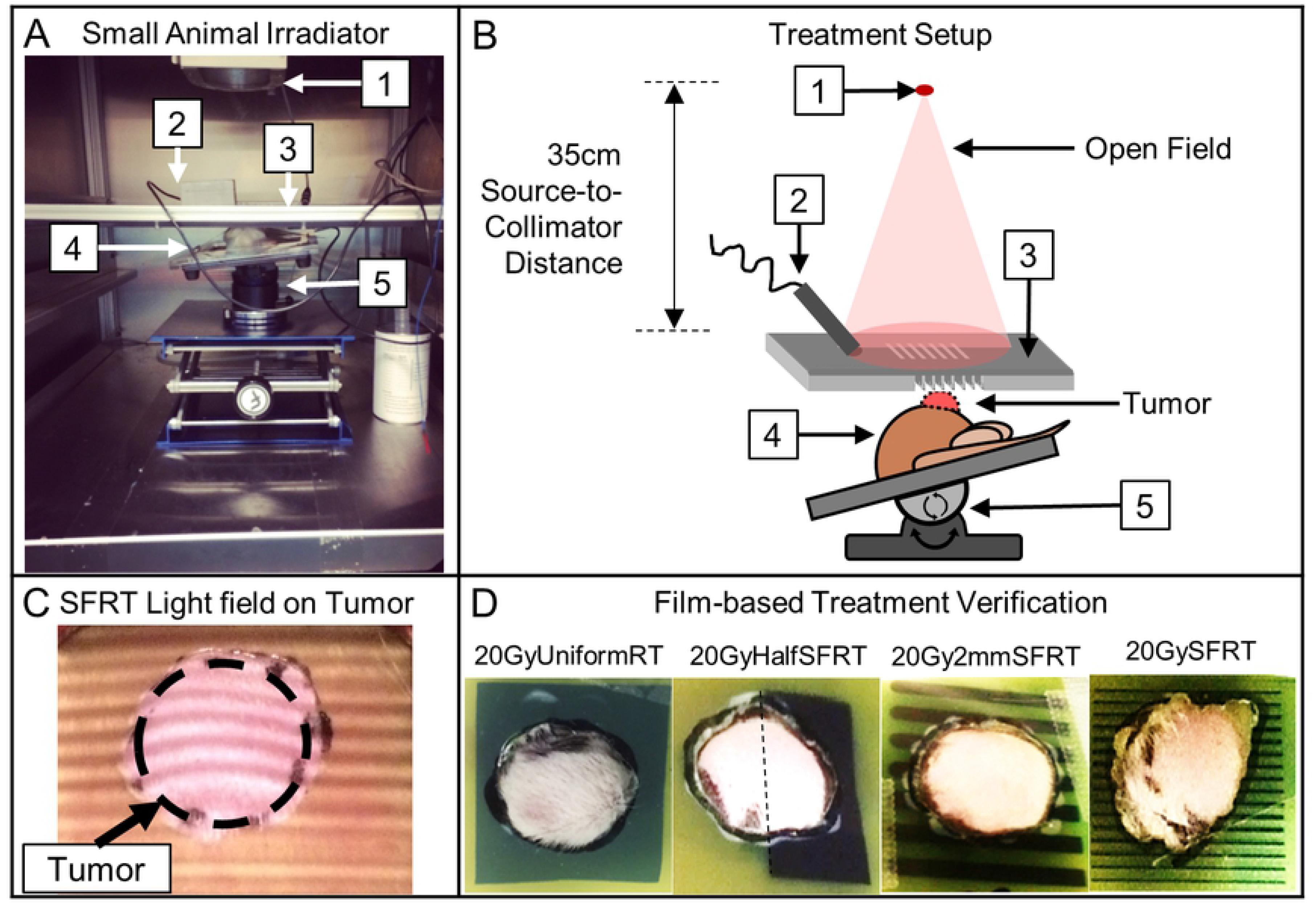
Animal irradiation setup and treatment alignment and verification. (A - B) The treatment setup components include (1) X-ray source, (2) endoscopic camera (lens shielded), (3) field shaping collimator for all treated arms (20GySFRT shown), (4) animal and tumor, and the (5) 3-axial heated animal positioning stage. (C) Photo of the built-in irradiator light shines through the 50GySFRT collimator and onto the outlined tumor as seen from the beams-eye view camera (live feed used to position tumor within the treatment fields.) (D) EBT-3 treatment verification films with a cutout in the tumor region. The films were reviewed for all treated animals for treatment targeting verification.

**Fig 3.**
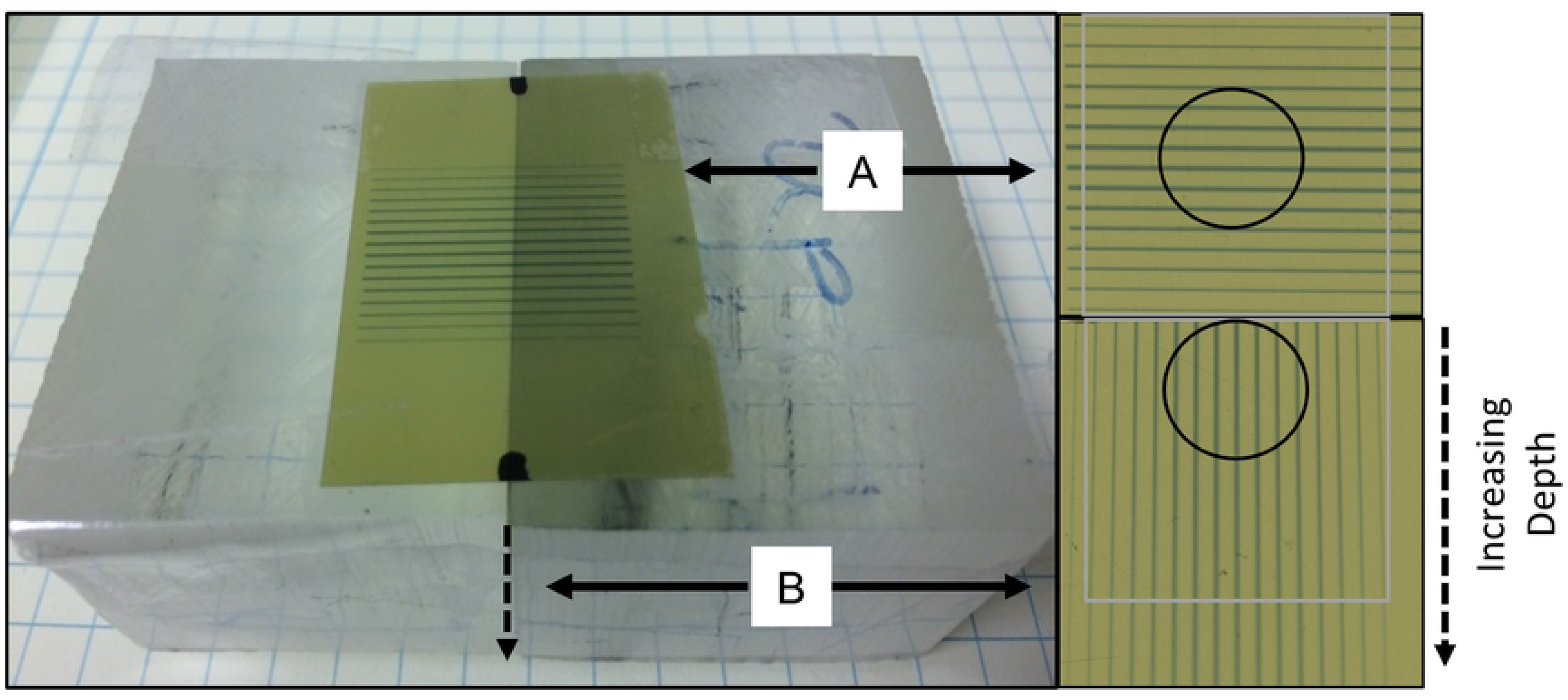
Phantom dosimetry measurement. EBT-3 films were calibrated by ion chamber under large field conditions. All beam profiles and corresponding percentage depth dose were measured using two films as shown: one is on the surface perpendicular to radiation beam (A) and one sandwiched between two small phantom blocks parallel to radiation beam (B). The circles indicate the film areas used for volume-average dose calculation estimates. The following assumption was made for volume-averaged tumor dose and EUD calculations: dose value does not vary +/−1cm along the direction parallel to the same valleys/peaks.

**Fig 4.**
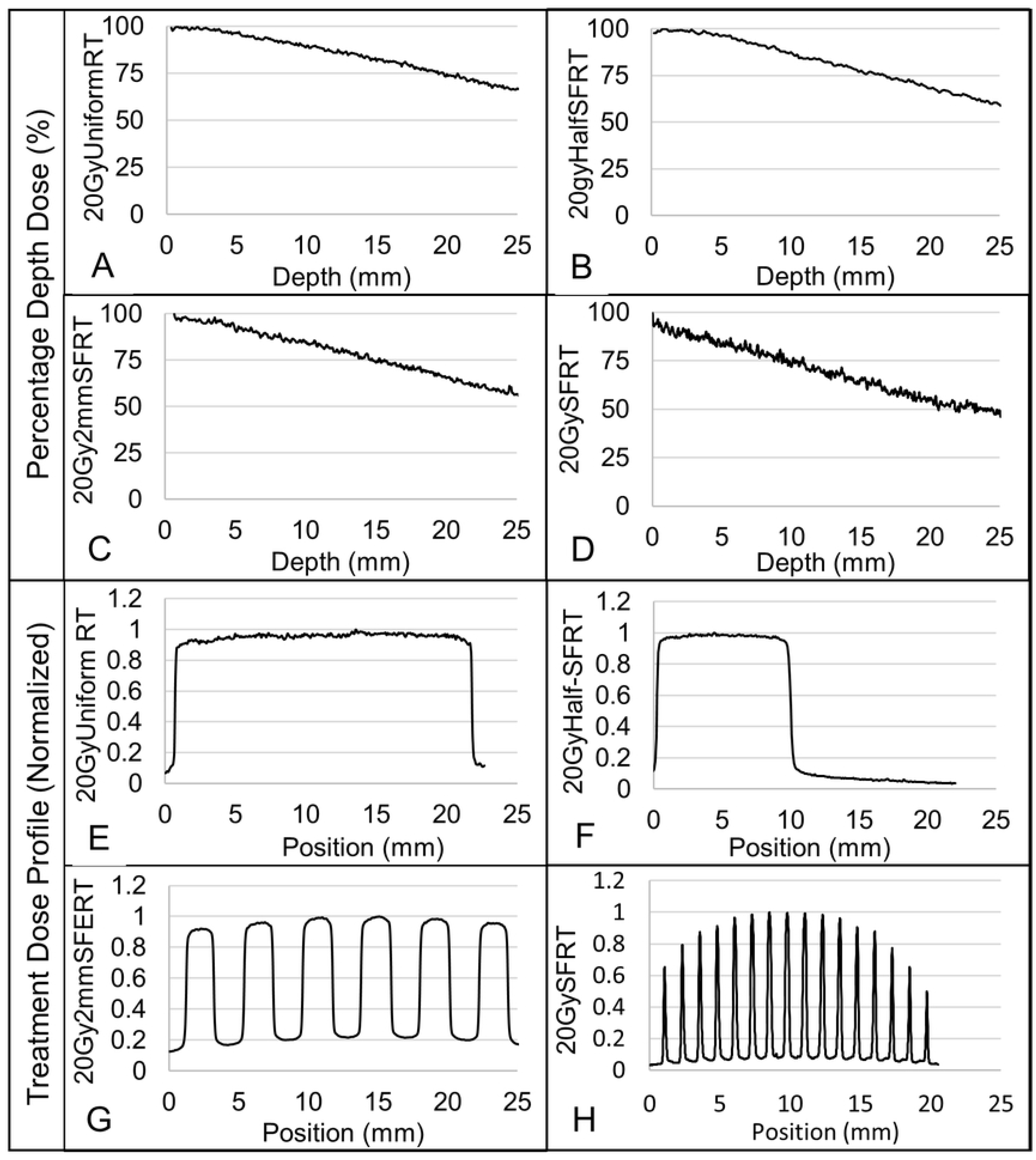
Measured dose beam profiles and percentage depth doses for all treatment arms. (A-D) Figures display the percentage depth doses for each of the 20Gy volume-averaged treatment arms. (E-H) Figures display the corresponding SFRT beam profiles for each of the 20Gy volume-averaged treatment arms. Note that the 20GySFRT and 50GySFRT arms share the same SFRT collimator and thus the same relative dosimetry. The large non-uniformity of the peak doses in the SFRT radiation is due to the finite x-ray target size and the nondivergence of the SFRT collimator. However, the actual peak dose non-uniformity in the treated tumor (diameter of ~10mm) is within 10%.

### Animal radiation delivery and verification

All of the RT collimators were aligned with x-ray target of the irradiator using film dosimetry. Animals were anesthetized with vaporized isoflurane mixed with oxygen carrier gas and positioned on an electronically controlled heating pad (Fig 2, panels A and B). For radiation tumor targeting we used the light field and a PC-linked camera before radiation and verified it with film dosimetry during each irradiation (Fig 2, panel C). Live video-feed from the camera was used for animal tumor-radiation alignment and for animal monitoring during treatment. Radiation targeting is achieved by (a) delineating the tumor boundary on animal skin using marker pre-treatment, (b) transferring the marking onto the verification EBT-3 film taped on skin and cutting out the tumor portion of the film, (c) taping the film back with the tumor inside the cutout, (d) placing the animal in the irradiator and align the tumor with the radiation, and (e) animal monitoring throughout irradiation. The treatment verification films were reviewed post-radiation for radiation targeting documentation (Fig 2, panel D).

### Tumor volume imaging and body weight monitoring

Three-dimensional B-mode ultrasound imaging of the tumors was performed using a Vevo 770 preclinical ultrasound scanner (Vevo 770, VisualSonics, Toronto, ON, Canada) and the resulting images used to calculate tumor volume, as described in a previous publication (*23*). Imaging was performed on the day before treatment as well as every third day post-treatment for approximately 30 days, or when maximal tumor burden was met, at which point the animals were humanely sacrificed per IACUC-approved animal protocol. Fig 5 shows an illustration of the 3D ultrasound tumor imaging setup and acquisition. Three-dimensional imaging is performed by mechanically stepping the ultrasound probe in the elevational dimension and acquired a two-dimensional image at each step (100um step size, 2cm elevational scan length). The reconstructed 3D ultrasound images were used to calculate tumor volume. The longest orthogonal tumor dimensions in each 3D image were measured using the digital caliper feature on the Vevo 770 imaging software and tumor volume was approximated using the volume formula for an ellipsoid, 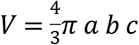, where V is the calculated tumor volume, and *a*, *b*, and c are each the half lengths of the principal axes of the tumor (*28*). A sample tumor volume change post radiation from a 20GyHalfSFRT arm animal shows no tumor control (Fig 5, panel D). Animal body weight was measured using the same schedule.

**Fig 5.**
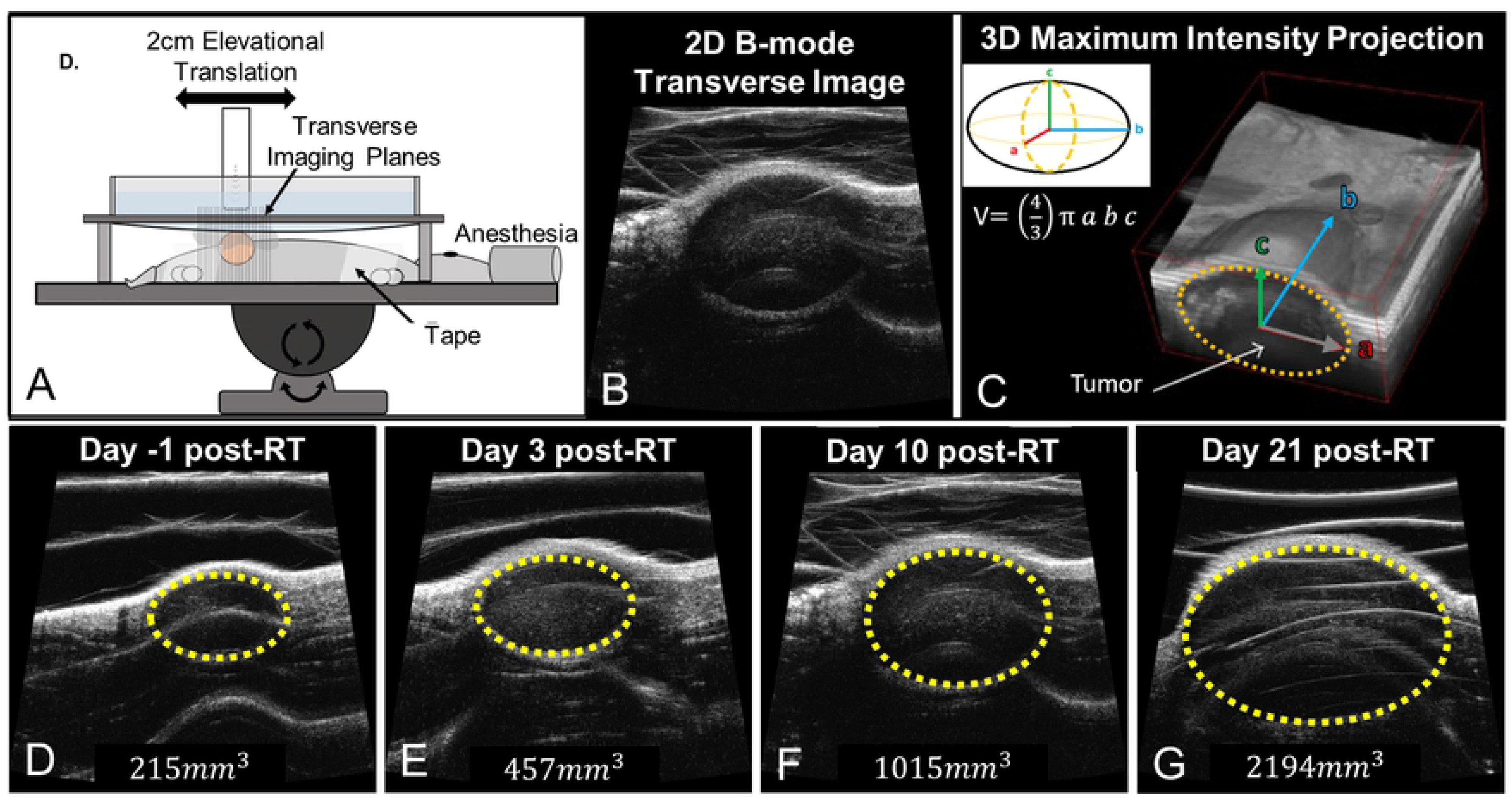
Illustration of 3D ultrasound imaging-based tumor volume measurement. Figure (A) is an illustration of the 3D ultrasound imaging setup with anesthetized animal (23). Two-dimensional transverse image slices (B) are acquired along the elevational direction and are then reconstructed into 3D images (29) (C). Tumors are visually identified on the ultrasound images. Resulting 3D images (C) are used to measure the tumor dimensions and calculate tumor volume. Imaging data is acquired pre-treatment (D) and every ~third day thereafter (E-G). In images D-G the tumor (yellow dotted line) and corresponding tumor volume grow over time following a 20GyHalfSFRT treatment.

### Association between SFRT dosimetry and treatment response

We analyzed the associations between animal treatment responses and each of the nine dosimetric parameters, listed in Table 1. The treatment responses are time-to-euthanasia, proportion of animals surviving to Day 17, and change in animal body weight on Day 17. We deem animal survival is a better indicator of tumor treatment response than tumor size change in this study. When tumors reach the maximum tumor mass, defined by the IACUC-approved animal protocol, ethical euthanasia is performed. As a result, animal numbers in different study arms decrease at different rates, which can introduce biases due to unbalanced sample sizes in the study. Hence, Day 17 was chosen for the linear regression association studies because at this timepoint there is a good compromise between the number of animals available for statistical consideration and the magnitude of radiation effects. We also fit a more robust Cox Proportional Hazards (CoxPH) model to the full data set that includes all animals.

Animal body weight change on Day 17 is used as an indicator of treatment toxicity. Animal body weight change is a gross assessment on treatment toxicity, especially in this study where tumors were implanted in the rodent flank, near the lower gastro-intestinal tract (including the rodent anus, rectum, colon, and cecum) and parts of the upper gastro-intestinal tract (including portions of the small bowel). We speculate that some treatment arms may induce more GI toxicity that others. We subtracted the tumor weight from the measured body weight and regard this “net” animal body weight change as an indication, not confirmation, of treatment toxicity. To confirm any lower GI toxicity, additional tissue histological staining or organ function examination studies would be necessary, both of which are beyond the scope of this work.

### Statistical methods

We computed Product-Limit (Kaplan-Meier) Estimator and Logrank (Mantel-Haenszel) test for statistical significance of survival difference between each pair of treatment arms (30). Multiple simple linear regression models (31) were used to study the association between dosimetric parameters with animal body weight and percentage survival within treatment group on Day 17. R^2^ (square of the Pearson correlation) coefficient is computed to estimate the proportion of variance explained in each of the linear regression models. In general, the greater the magnitude of the test statistic (t or F), the more closely associated the dosimetric parameter studied is with the treatment response (survival or body weight).

In addition to linear regressions, we fit Cox Proportional Hazard (CoxPH) models with individual animal survival as the time-to-event outcome, which used data from all dates including Day 17. This allowed us to calculate the hazard ratio associated with the impact of dosimetric parameters on treatment response. We also used a Pearson Correlation matrix to show the cross-correlation between each pair of the dosimetric parameters. All data collected were analyzed using R (version 3.5.3) statistical software available from R Core Team.

## Results

### Overall treatment response

Fig 6 shows (A) animal survival, (B) normalized tumor volume, and (C) normalized body weight post treatment for all 6 study arms. In this study no animal died of body condition deterioration. All endpoints were due to ethical animal euthanasia triggered by tumors exceeding the maximum allowable burden per IACUC-approved animal protocol limitations. Our data shows that the 20GyUnformRT arm has the best tumor control followed by the 50GySFRT and 20Gy2mmSFRT arms. Note that among the four arms sharing similar volume-averaged dose (20Gy or 18Gy) survival varies greatly, from 33% to 100% on Day 17, which is a strong indication that volume-averaged dose is poorly associated with tumor treatment response. The tumor volume data indicate that although 50GySFRT arm and 20Gy2mmSFRT arm have similar survival the former has a better tumor volume reduction than the latter arm. Only the 20GyUniformRT arm experienced weight loss post-treatment and then recovered back to pre-treatment weight after week three. The 20GySFRT and 20Gy2mmSFRT arms experienced similar body weight gains as the untreated arm, indicating little treatment toxicity from the two SFRT treatments.

**Fig 6.**
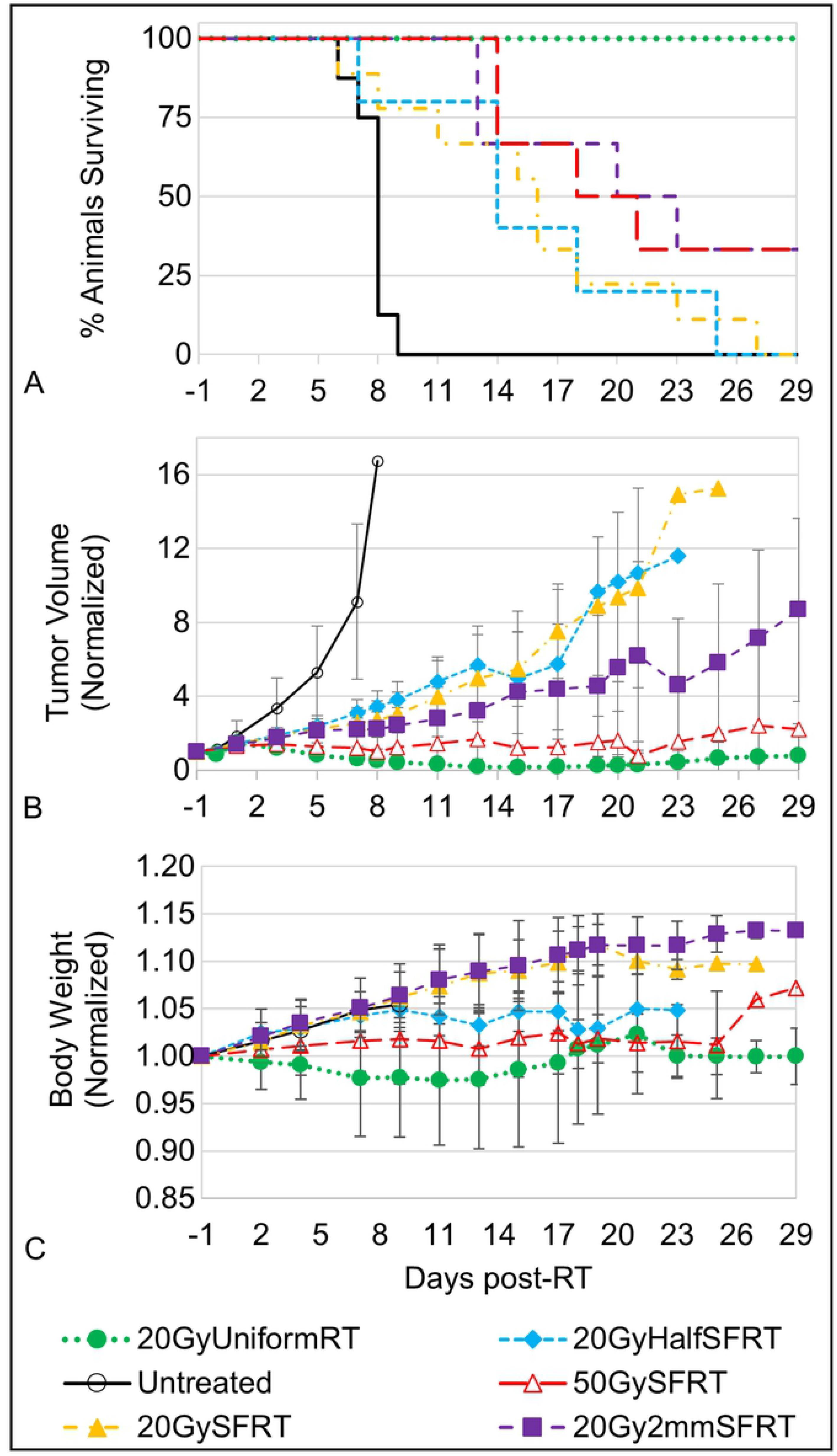
Animal survival, tumor volume change, and body weight change post-treatment. Animal survival (A), normalized tumor volume (B), and normalized body weight (C) are shown for all six study arms. The differences between survival curve pairs are significant (p<0.05) for 20GyUniformRT-50GySFRT, 20GyUniformRT-20GyHalfSFRT, 20GyUniformRT-20Gy2mmSFRT, 20GyUniformRT-Untreated, 20GyUniformRT-20GySFRT, Untreated– 20GySFRT, Untreated-20Gy2mmSFRT, Untreated-50GySFRT, and Untreated-20GySFRT, and moderately significant (0.1>p<0.05) for 20GyHalfSFRT-50GySFRT, 20Gy2mmSFRT-50GySFRT, and 20GySFRT–50GySFRT.

### Association between tumor response and SFRT dosimetry

We associated eight dosimetric parameters with percentage of animals surviving to Day 17 and with the survival curves shown in Fig 6. Fig 7 shows scatter plots of eight tumor-related dosimetric parameters vs. percentage survival at Day 17, each fitted with a corresponding regression line, R^2^ (Fig 7). Tumor EUD (R^2^=0.7923, F-stat=15.26*), Valley dose (R^2^=0.7636, F-stat=12.92*), and percentage volume directly irradiated (R^2^=0.7153, F-stat=10.05*) are the top three most statistically significant dosimetric parameters in terms of association with the animal survival at Day 17 (see Table S1). Peak dose (R^2^=0.04472, F-stat=0.6874 (not sig.)) and AVG Dose (R^2^ = 0.2745, F-stat=1.514 (not sig.)) showed little association with survival.

**Fig 7.**
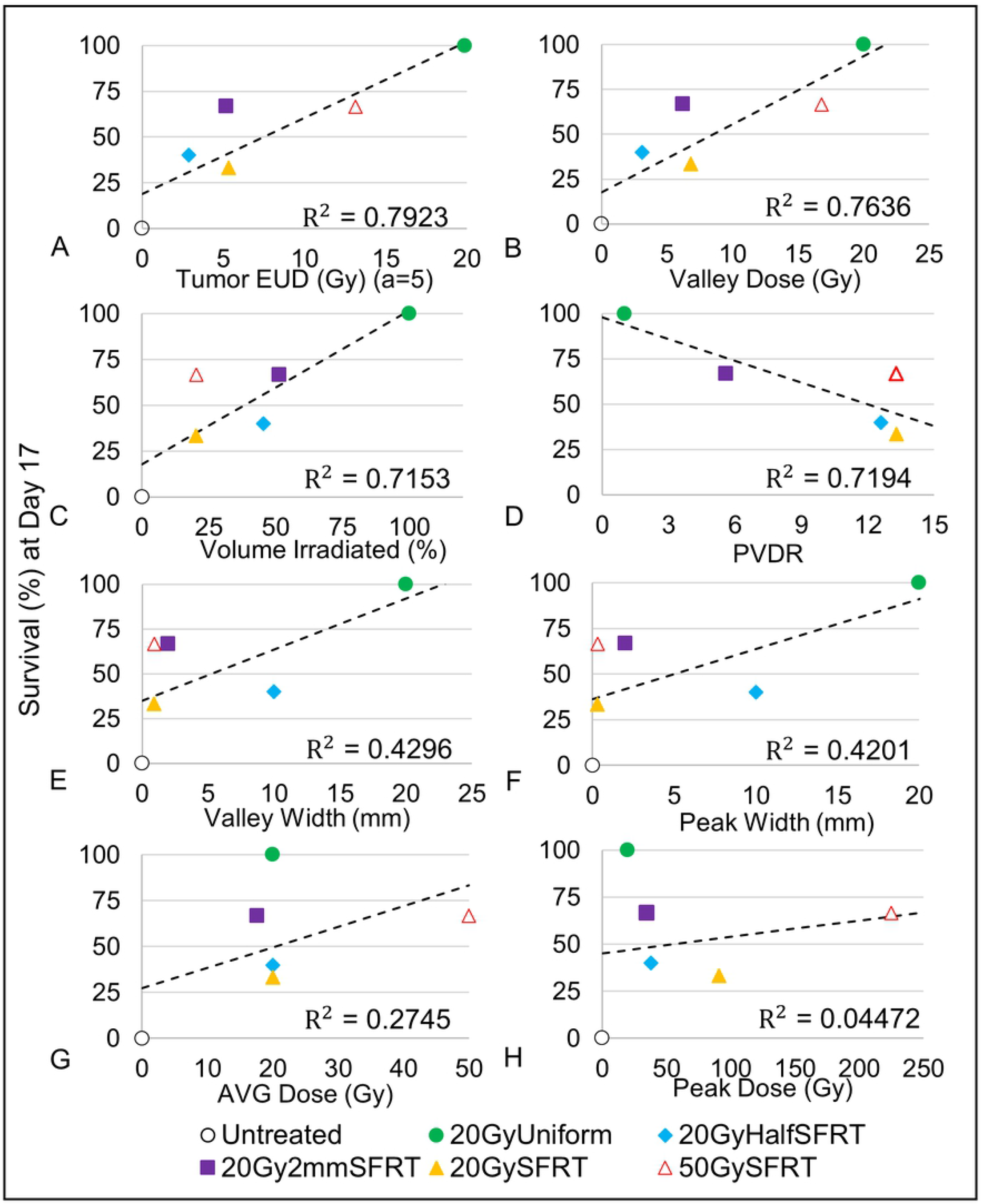
Associations between Percentage Survival (Day 17) and eight dosimetric parameters. Tumor EUD (A), valley dose (B), percentage volume irradiated (C), valley width (D), peak width (E), volume-averaged dose (F), peak dose (G), and PVDR (H) vs survival (%) at Day 17 are presented as well as their corresponding regression lines and R^2^ values. Eight linear regression models with single covariates, one for each dosimetric parameter, were used to calculate the R^2^ value and corresponding statistics.

To validate the above finding in Fig 7 we used data from the entire survival curves in Univariate Cox Proportional Hazards analysis and the results are shown in Table 2. The results from the Univariate Cox Proportional Hazards analysis confirms the results from the linear regression analysis - among the eight dosimetric parameters analyzed tumor EUD (z-stat=−4.07***), valley/min dose (z-stat=−4.338***), and percentage tumor volume directly irradiated (z-stat=−3.837***) have the closest associations with animal survival. Compared to the linear regression analysis (Fig 7) the improved p-values in the CoxPH model analysis is likely due to the increased sample size. The Hazard Ratio shows the impact of change in each of the dosimetric parameters to the hazard rate (risk of death). For instance, when valley/min dose parameter changes by 1 Gy, the hazard rate (risk of death) changes by 19% (95% CI, 26% - 11%) with p-value of 1.44×10^−5^. For a 1Gy change in peak dose, the corresponding change in hazard rate is 0.2% (95% CI, 0.7% - 0.3%) with p-value of 0.432. Three additional statistical tests were used to validate the CoxPH z-test statistics results for each model (Likelihood Ratio Test, Wald Test, and Logrank Test) and all three tests largely agree with the results presented in Table 2.

**Table 2:**
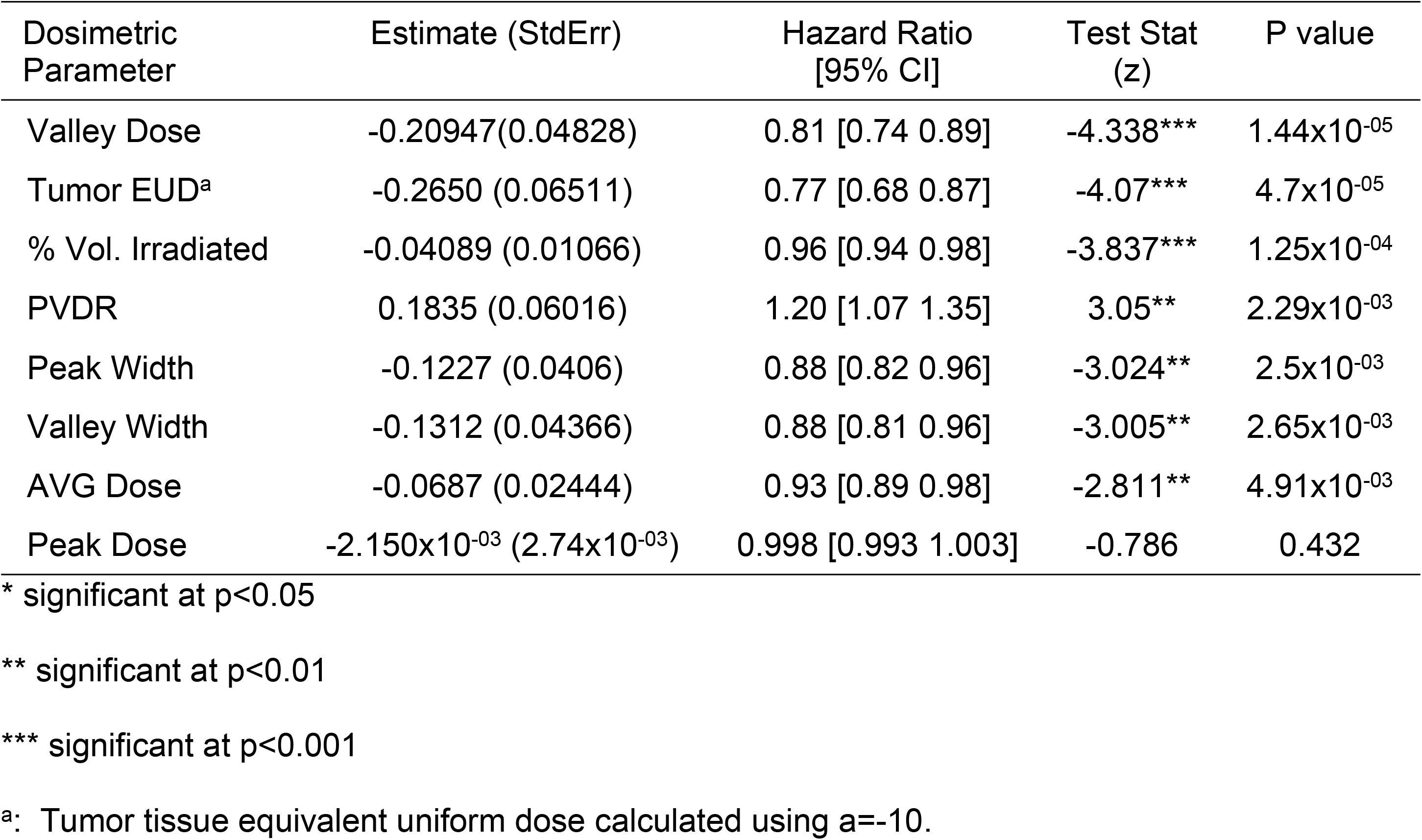
Table of coefficients for univariate Cox Proportional Hazards analysis of survival.

| Dosimetric Parameter | Estimate (StdErr) | Hazard Ratio [95% CI] | Test Stat (z) | P value |
| --- | --- | --- | --- | --- |
| Valley Dose | -0.20947(0.04828) | 0.81 [0.74 0.89] | -4.338*** | $1.44 \times 10^{-5}$ |
| Tumor EUD<sup>a</sup> | -0.2650 (0.06511) | 0.77 [0.68 0.87] | -4.07*** | $4.7 \times 10^{-5}$ |
| % Vol. Irradiated | -0.04089 (0.01066) | 0.96 [0.94 0.98] | -3.837*** | $1.25 \times 10^{-4}$ |
| PVDR | 0.1835 (0.06016) | 1.20 [1.07 1.35] | 3.05** | $2.29 \times 10^{-3}$ |
| Peak Width | -0.1227 (0.0406) | 0.88 [0.82 0.96] | -3.024** | $2.5 \times 10^{-3}$ |
| Valley Width | -0.1312 (0.04366) | 0.88 [0.81 0.96] | -3.005** | $2.65 \times 10^{-3}$ |
| AVG Dose | -0.0687 (0.02444) | 0.93 [0.89 0.98] | -2.811** | $4.91 \times 10^{-3}$ |
| Peak Dose | $-2.150 \times 10^{-3}$ ($2.74 \times 10^{-3}$) | 0.998 [0.993 1.003] | -0.786 | 0.432 |

\* significant at $p < 0.05$
\*\* significant at $p < 0.01$
\*\*\* significant at $p < 0.001$
<sup>a</sup>: Tumor tissue equivalent uniform dose calculated using $\alpha = -10$ .

### Association between body weight change and SFRT dosimetry

Eight dosimetric parameters are associated with the body weight change on Day-17. Note that the body weight is the measured body weight subtracted the measured tumor weight to remove the influence of tumor size on the analysis. Fig 8 is a scatter plot of the dosimetric parameters vs. the “net” body weight at Day 17. This time point was chosen for both the tumor and body weight study because it is a good compromise between data statistics and magnitude of treatment response. Table 3 is a table of coefficients for the corresponding linear regression models used in Fig 8. In general, the greater the magnitude of the t statistic, the greater the individual parameter association with Body Weight (Day 17). For the F-statistic, the greater the statistic value, the more closely associated the model is with Body Weight (Day 17). Based on the t statistics and F-statistics, among the eight dosimetric parameters studied the Valley Dose has the greatest, yet modest, association with Body Weight (Day 17). The order of decreasing association with the body weight change are: valley dose (R^2^=0.3814, F-stat=13.45**), valley width (R^2^=0.2853, F-stat=8.783*), peak width (R^2^=0.2759, F-stat=8.382*), percentage volume irradiated (R^2^=0.1985, F-stat=5.448*), PVDR (R^2^=0.1203, F-stat=3.009 (not sig.)), volume-averaged dose (R^2^=0.03308, F-stat=0.7526 (not sig.)), normal tissue EUD (R^2^=1.022×10^−03^, F-stat=0.882 (not sig.)), and peak dose (R^2^=5.99×10^−06^, F-stat=1.32×10^−04^ (not sig.)). A strong similarity between the peak width and valley width association with body weight is expected (see discussion in 4E section). No significant association is observed between body weight change post radiation and PVDR, average dose, normal tissue EUD, and peak dose.

**Table 3:** Table of coefficients for univariate linear regression analysis of Body Weight (Day 17)

| Dosimetric Parameter | Estimate (StdErr) | R<sup>2</sup> | t value | F-statistic |
| --- | --- | --- | --- | --- |
| Valley Dose | -6.306x10<sup>-03</sup> (1.713x10<sup>-03</sup>) | 0.3814 | -3.683** | 13.56** |
| Valley Width | -4.498x10<sup>-03</sup> (1.518x10<sup>-03</sup>) | 0.2853 | -2.964** | 8.783** |
| Peak Width | -4.333x10<sup>-03</sup> (1.497x10<sup>-03</sup>) | 0.2759 | -2.895** | 8.382** |
| %Vol. Irradiated | -9.519x10<sup>-04</sup> (4.08x10<sup>-04</sup>) | 0.1985 | -2.334* | 5.448* |
| PVDR | 4.525x10<sup>-03</sup> (2.609x10<sup>-03</sup>) | 0.1203 | 1.735 | 3.009 |
| AVG Dose | -1.054x10<sup>-03</sup> (1.215x10<sup>-03</sup>) | 0.03308 | -0.867 | 0.7526 |
| Tissue EUD<sup>a</sup> | -6.184x10<sup>-05</sup> (4.123x10<sup>-04</sup>) | 1.022x10<sup>-03</sup> | -0.15 | 0.882 |
| Peak Dose | -2.252x10<sup>-06</sup> (1.961x10<sup>-04</sup>) | 5.99x10<sup>-06</sup> | -0.011 | 1.32x10<sup>-04</sup> |

\* significant at p<0.05
\*\* significant at p<0.01
<sup>a</sup>: Normal tissue equivalent uniform dose calculated using $\alpha=5$ .

**Fig 8.**
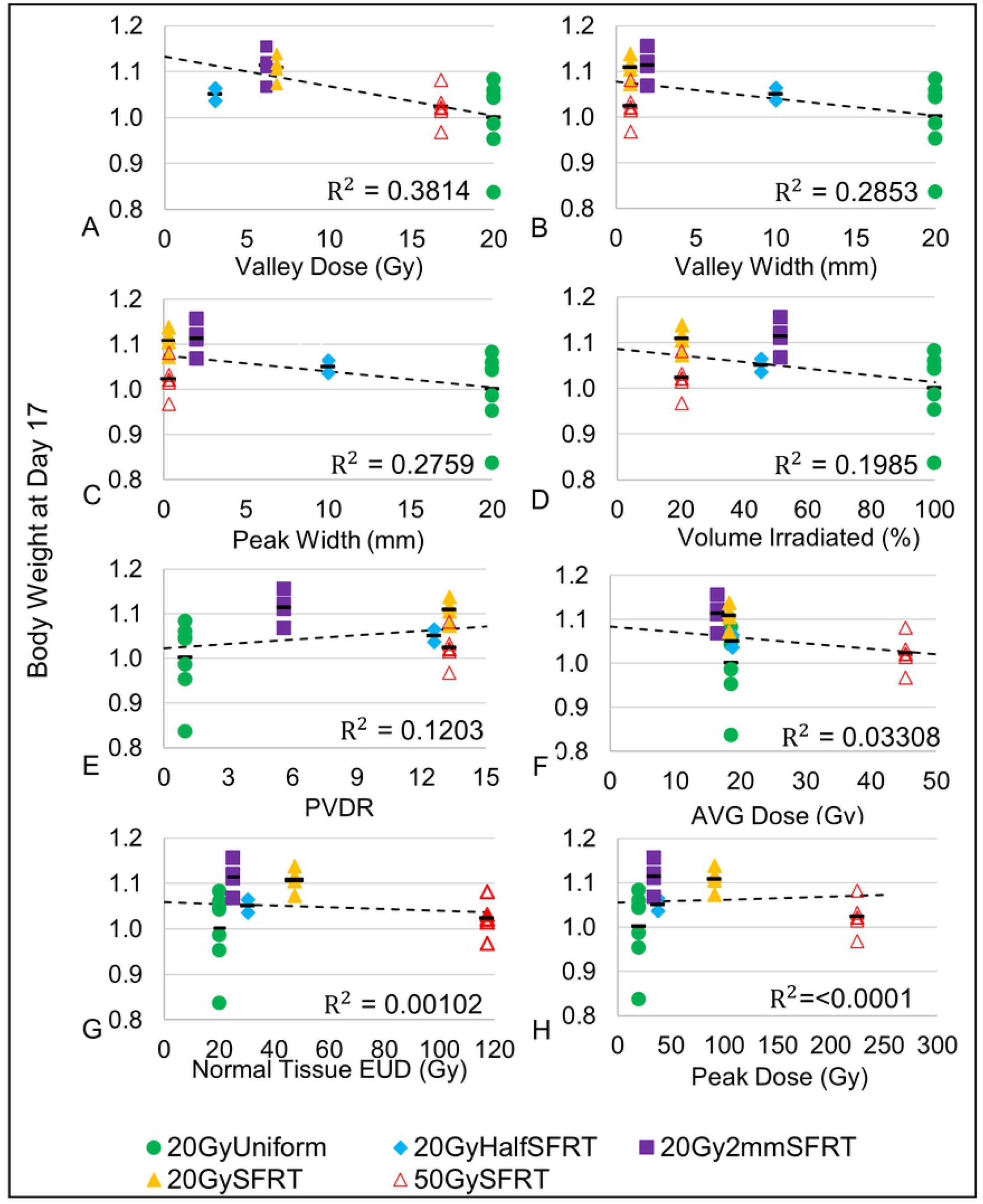
Associations between Body Weight (Day 17) and eight dosimetric parameters. Scatter plots of each of the 8 treatment dosimetric parameters: valley dose (A), valley width (B), peak width (C), percentage volume irradiated (D), normal tissue EUD (E), PVDR (F), volume-averaged dose (G), and peak dose (H) vs % Body Weight at Day 17 and their corresponding regression lines and R^2^ values are shown. Eight linear regression models with single covariates, one for each dosimetric parameter, were used to calculate the R^2^ value and corresponding statistics.

## Discussion

### Study limitations

There are several limitations in this study, many of which are discussed below. (i) There was no image-guidance used in the irradiation study. Our remedy for the lack of online imaging technology included the use of light field and video-based animal alignment, of treatment verification film, and lastly, removal of treatment-misaligned animals from the study. This was judged from reviewing the treatment verification film for each animal. Our remedy worked well resulting in a 20GyHalfRT arm % volume irradiated of 47.8.8% (±2.2) near the target value of 50% (S1 Fig1) (ii) No CT-based treatment planning. Based on the anatomical location of the implanted tumor (rodent flank) we believe a portion of the rodent GI tract may have been irradiated but the actual volume irradiated is unknown. Because all animals were randomized across study arms such that all arms have the same average pre-treatment tumor size and similar tumor location distribution, it is reasonable to assume that any variations in portions of GI track irradiated do not bias any particular study arm. (iii) Only a single tumor model used. The FSA rat tumor model does not represent tumors with low vascularity, which may have different treatment responses. The study should be repeated using different tumor and animal models. (iv) The dosimetric parameters have strong cross correlations in this study, which is discussed in more detail at the end of this section.

The potential impact of spatial fractionation pattern (lines vs. dots, for instance) on treatment response is beyond the scope of this work. However, it is a very important question that deserves methodical investigations as some spatial fractionation patterns are easier to achieve than others in practical application. Our data shows that valley/minimum dose has the closest association with treatment response for tumor and body weight. However, different spatial fractionation patterns with the same valley dose may not lead to the same treatment response when a different endpoint is used. In our study the 20Gy2mmSFRT arm and the 20GySFRT arm have similar valley doses but dissimilar survival fraction on Day 17. To investigate the impact of radiation spatial fractionation pattern alone on given treatment responses, carefully designed new studies are needed.

The exciting noncytotoxic effects of SFRT, such as induction therapy to sensitize tumor to increase therapeutic ratio of the following therapy including anti-tumor immunotherapy, remain largely underexplored (32); however, they are also beyond of the scope of this work. Our own and others’ work have demonstrated that SFRT radiation impacts tumor microenvironment and modulates immune system very differently than uniform radiation therapy (33–35). We intend to conduct similar studies to identify associations between dosimetric parameters and these indirect effects of SFRT in the future.

### SFRT dosimetric association with treatment tumor response

#### Valley dose and tumor EUD

The importance of tumor minimum dose to tumor control has long been established in conventional radiation therapy (36). Does the same association between tumor control and minimum/valley dose hold for SFRT? For some the answer is yes and sophisticated techniques have been developed to “fill up” the dose valleys in an MRT beam by interlacing the microbeams from MRT from different irradiation angles. As a result, a uniform dose distribution inside the tumor is reached (*37*) while the surrounding normal tissue out of the “cross-firing” range still receive largely MRT radiation pattern of peaks and valleys. In a synchrotron-MRT study Ibahim et al. (38) reported that valley dose is closely correlated with cell survival, but valley dose alone does not determine the observed radiobiological effects. Our study shows that the tumor EUD (a=−10) and minimum/valley tumor dose have the highest linear associations (R^2^=0.7923, F-stat=15.26*; R^2^=0.7636, F-stat=12.92*, respectively) with tumor treatment response (Fig 7 and S1 Table). This observed association between tumor treatment response with tumor valley/minimum dose and tumor EUD dose in this preclinical study is consistent with their known association in tumor treatment response seen in clinical conventional uniform dose radiation therapy.

Our data suggests that valley/minimum dose or Tumor EUD are more appropriate than peak dose for SFRT treatment prescription. When tumor control is the endpoint, we suggest that equal valley or minimum dose be used for comparative study between a uniform radiation and SFRT therapy or among different SFRT treatments.

#### PVDR

Our data showed that PVDR has a consistent but not statistically significant association with tumor treatment response (R^2^=0.7194, F-stat=7.691) (Fig 7, S1 Table). The linear regression analysis on day 17 was not statistically significant. The CoxPH analysis using the entire survival data set show a modest association with survival. Although not statistically significant, an inverse association is observed between PVDR value and survival fraction on Day 17 - the higher PVDR value the less survival fraction. The inverse association is largely determined by the uniform radiation arm where PVDR value is 1.0. If this data point is removed, the PVDR association with survival for all SFRT arms is inconclusive (Fig. 7). We believe this result of inverse association is likely biased by the study design that has very limited PVDR values (4 values) and strong cross-correlations between PVDR and other SFRT parameters (see more discussion later in the section). A better understanding of PVDR’s association with a given treatment response requires a carefully designed new study that focuses on the impact of PVDR value on treatment response.

#### Percentage volume irradiated, peak width, and valley width

It seems logical that tumor treatment response is closely associated with the tumor volume irradiated. However, this is not supported by a clinical GRID therapy study by Neuner et al. (2) where both MLC-based and collimator-based GRID treatments showed similar response rates for pain, mass effect, other patient complaints, and have similar adverse reactions. The collimator-generated GRID had 50% of the radiation field open while the MLC-generated GRID had only 31% open. In our study the 20GyHalfSFRT and 20Gy2mmSFRT arms have similar percentage-volume-irradiated (as well as PDD curves) but there is a difference of 5 days in the 50% survival time (Fig 1 and Fig 6). Nonetheless, our data shows that percentage-volume-irradiated has the 3rd highest linear association (R^2^ = 0.7153, F-stat=10.05*) with tumor treatment response (Fig 7 and Table 2). Since percentage-volume-irradiated is jointly determined by peak width and valley width it is understandable to see moderate associations between tumor treatment response and peak width (R^2^=0.4201, F-stat=2.898 (not sig.)) and valley width (R^2^=0.4296, F-stat=3.012 (not sig.)). In a synchrotron microbeam brain study using multiple beams Serduc et al. kept valley dose constant while varying peak width and peak dose. They concluded that the latter two parameters have strong influence therapeutic ratio (39).

#### Volume-averaged dose and peak dose

This study is designed to scrutinize the association of volume-averaged dose with tumor treatment response (Fig 1). The four study arms sharing very similar volume-averaged doses (20 or 18 Gy) exhibited very different tumor treatment responses (Fig 6 and 7) showing the survival rate at day 17 varied from 100% to 33%. Therefore, the association between volume-average dose and tumor treatment response is weak.

We found that peak dose has little to no association with tumor treatment response (R^2^=0.04472, F-stat=0.6874 (not sig.)) (Fig 7, S1 Table, Table 2). This finding is significant because peak dose has been used for treatment prescription in practically all SFRT treatments (*8*) (*9*). Although the linear regression analysis on day 17 showed a weak association between peak dose and survival that was not statistically significant, the CoxPH analysis using the entire survival data set did show a modest association with survival.

### SFRT dosimetric association with normal tissue toxicity

We did not study treatment induced normal tissue toxicity directly in this study. We used body weight change post radiation (targeted to the flank, lower abdominal region of the animal) as an indicator, not an evidence of normal tissue toxicity. We did not see a strong association between animal body weight change and any of the eight dosimetric parameters studied, except a modest association with valley/minimum dose.

#### Valley dose

The strongest association we observed is a weak one between body weight change and valley/min dose (R^2^=0.3814, F-stat=13.56**) (Table 3). Note that valley/min dose is also strongly associated with tumor treatment response (R^2^=0.7636, F-stat=12.92*). Our finding is consistent with a normal mouse brain MRT study Nakayma et al. reported that valley dose is one of the important factors to determine normal brain dose tolerance (40). Our data suggests that valley dose may have a close correlation with both tumor control and toxicity, and thus is a crucial dosimetric parameter in SFRT treatment.

#### Valley width, peak width, percentage volume irradiated

The valley width, peak width, and percentage volume of the tumor that is irradiated parameters were only weakly associated with animal body weight change post radiation (R^2^=0.2853, F-stat=8.783**; R^2^=0.2759, F-stat=8.382**; and R^2^=0.1985, F-stat=5.448*, respectively) (Fig 8 and Table 3). Note that in this study peak width and valley width are closely correlated (more discussion on correlations, below). Percentage volume directly irradiated showed no statistically important association with body weight change. Neuner et al. reported that they observed similar treatment responses from clinical GRID treatments of different percentages of volume directly irradiated (*2*).

#### Normal tissue EUD, PVDR, volume-averaged dose, peak dose

The normal tissue EUD, PVDR, volume-averaged dose, and peak dose parameters showed little to no association with body weight change post radiation (R^2^=1.022×10^−03^, F-stat=0.882 (not sig.); R^2^=0.1203, F-stat=3.009 (not sig.); R^2^=0.03308, F-stat=.7526 (not sig.); and R^2^=5.99×10^−06^, F-stat=1.32×10^−04^ (not sig.), respectively). Our finding is consistent with a rat normal brain minibeam study by Prezado et al. showing arms with similar volume-average-doses have drastic differences in survival (*14*) and inconsistent with a MRT study on normal mouse skin by Priyadarshika et al. concluded that integrated dose (i.e., volume-averaged dose) rather than peak or valley dose, may dictate the acute skin toxicity (41).

#### 2mm wide beam array SFRT

Our data indicates that the 20Gy2mmSFRT arm is not only the most relevant to clinical application because of its millimeter scale, but it also has the potential for superior therapeutic ratio. The 20Gy2mmSFRT arm showed similar survival with the 50GySFRT arm but has significantly lower valley dose (6.2 Gy vs. 17 Gy). At the same time, it showed the least, if any, body weight change compared to the untreated arm while the 50GySFRT arm with 0.31mm beam width exhibited significant body weight growth deficit (Table 1 and Fig 6). The 20GyUniform arm has the best tumor treatment response and the worst body weight change. Our data indicated the 2mm wide beam array is a kV photon SFRT pattern that has the potential for high therapeutic ratio SFRT and deserves further investigation.

### Cross-correlation in the SFRT dosimetry parameters

The dosimetric parameters studied in this work are not all independent variables and their cross-correlations are shown in the table of Pearson Correlation coefficients (Table 4.) The larger the magnitude of the coefficient, the more co-linear and correlated the pair of dosimetric parameters. In this study, peak width and valley width are perfectly co-linear (correlation of 1.0) by study design. Valley/min dose, a parameter used in tumor EUD calculation, is also highly correlated with tumor EUD (correlation of 0.99). These strong correlations explain the similar statistical associations of these parameters with treatment responses. These correlations also limited the study’s ability to better exam the association between a given treatment response with each of the dosimetric parameters.

**Table 4:** Pearson Correlation coefficient matrix for the eight SFRT dosimetric parameters relevant for tumor treatment response.

|  | Valley<br>Dose | Peak<br>Dose | AVG<br>Dose | Tissue<br>EUD | Tumor<br>EUD | Peak<br>Width | Valley<br>Width | PVDR | % Vol.<br>Irradiated |
| --- | --- | --- | --- | --- | --- | --- | --- | --- | --- |
| Valley<br>Dose | 1.00 |  |  |  |  |  |  |  |  |
| Peak<br>Dose | 0.38 | 1.00 |  |  |  |  |  |  |  |
| AVG<br>Dose | 0.63 | 0.91 | 1.00 |  |  |  |  |  |  |
| Tissue EUD | 0.44 | 0.99 | 0.95 | 1.00 |  |  |  |  |  |
| Tumor EUD | 0.99 | 0.26 | 0.53 | 0.32 | 1.00 |  |  |  |  |
| Peak Width | 0.63 | -0.36 | 0.00 | -0.28 | 0.72 | 1.00 |  |  |  |
| Valley Width | 0.65 | -0.34 | 0.02 | -0.25 | 0.74 | 1.00 | 1.00 |  |  |
| PVDR | -0.57 | 0.64 | 0.40 | 0.61 | -0.67 | -0.78 | -0.77 | 1.00 |  |
| % Vol. Irradiated | 0.70 | -0.26 | 0.13 | -0.17 | 0.78 | 0.93 | 0.93 | -0.94 | 1.00 |

## Summary

In this conventional dose rate small animal SFRT study we used a large range of radiation spatial fractionation scales to study the association of dosimetric parameters with treatment response. We concluded that valley/minimum dose, tumor EUD, and percentage tumor irradiated have strong and proportional associations with tumor treatment response while peak dose exhibited little association. Among the SFRT dosimetric parameters studied valley/min dose also showed the highest but modest association with body weight change post radiation.

## Acknowledgments

One of the authors (Chang) acknowledges Dr. Mark W. Dewhirst for his decade long unwavering encouragements and expert radiobiology advices, which are invaluable for this (and other) original work exploring the magic of radiation spatial fractionation. One of authors (Rivera) expresses her appreciation to Leith Rankine, MS for his kind help on EUD calculation.

## Supporting Information

**S1 Fig. Illustration of treatment delivery verification analysis by film.**(A) The post-treatment verification film for a 20GyHalfSFRT treated tumor shows that only one-half the tumor was treated as intended. The black dashed line in the photograph was drawn to illustrate which half of the tumor was irradiated. (B) The verification films for all 5 animals included in the study arm were analyzed by calculating the percentage area of the tumor irradiated.

**S1 Table. Univariate linear regression analysis of Survival Day 17.**This is the full table of coefficients for the corresponding linear regression models used in Fig 7. We analyze 8 models with single covariates, one for each dosimetric parameter and list their corresponding statistics. Tumor EUD and Valley Dose have the largest magnitude of effect on Survival (Day 17) and together with % Volume Irradiated are statistically significant. However, analyzing data for a single timepoint (Day17) is limited by animal losses at Day 17 (missing data), so we include a more robust statistical model that utilizes all the data in **Table 3**.

